# Targeted sequence capture of *Orientia tsutsugamushi* DNA from chiggers and humans

**DOI:** 10.1101/2021.01.07.425812

**Authors:** Ivo Elliott, Neeranuch Thangnimitchok, Mariateresa de Cesare, Piyada Linsuwanon, Daniel H. Paris, Nicholas PJ Day, Paul N. Newton, Rory Bowden, Elizabeth M. Batty

## Abstract

Scrub typhus is a febrile disease caused by *Orientia tsutsugamushi*, transmitted by larval stage Trombiculid mites (chiggers), whose primary hosts are small mammals. The phylogenomics of *O. tsutsugamushi* in chiggers, small mammals and humans remains poorly understood. To combat the limitations imposed by the low relative quantities of pathogen DNA in typical *O. tsutsugamushi* clinical and ecological samples, along with the technical, safety and cost limitations of cell culture, a novel probe-based target enrichment sequencing protocol was developed. The method was designed to capture variation among conserved genes and facilitate phylogenomic analysis at the scale of population samples. A whole-genome amplification step was incorporated to enhance the efficiency of sequencing by reducing duplication rates. This resulted in on-target capture rates of up to 93% for a diverse set of human, chigger, and rodent samples, with the greatest success rate in samples with real-time PCR C_t_ values below 35. Analysis of the best-performing samples revealed phylogeographic clustering at local, provincial and international scales. Applying the methodology to a comprehensive set of samples could yield a more complete understanding of the ecology, genomic evolution and population structure of *O. tsutsugamushi* and other similarly challenging organisms, with potential benefits in the development of diagnostic tests and vaccines.

## Introduction

Scrub typhus is a vector-borne zoonotic disease risking life-threatening febrile infection in humans. The disease is caused by an obligate intracellular Gram-negative bacterium, *Orientia tsutsugamushi.* Scrub typhus has an expanding known distribution, with most disease occurring across South and East Asia and parts of the Pacific Rim.

The genus *Orientia* is classified in the family Rickettsiaceae, a member of the order Rickettsiales. Two species of *Orientia* are currently recognised - *O. tsutsugamushi* and *O. chuto*, the latter known solely from a patient infected in the United Arab Emirates ^1^. Recent molecular identification of *O. tsutsugamushi* in humans in Chile ^2^ and 16S sequences with close homology to *O. tsutsugamushi* in dogs in South Africa ^3^ and small mammals in Senegal and France ^4^, and to *O. chuto* in chiggers in Kenya ^5^, suggest the possibility of further species and future taxonomic re-evaluation.

Larval trombiculid mites (chiggers) transmit *Orientia* to vertebrates, including man. The organism appears to be maintained by transovarial (vertical) and transstadial (between life-stages) transmission in chiggers, suggesting that they act as both vector and reservoir ^6–9^. There is good evidence for the transmission of *O. tsutsugamushi* to man by at least 10 species of chiggers ^10^. The ecology of the disease and the interaction of *Orientia* between vectors, small mammals and humans are complex and relatively poorly understood ^11^.

A high degree of phenotypic and genotypic diversity has been reported in *O. tsutsugamushi*. Several antigenic types appear to be widely present throughout Southeast Asia, with one (TA716) making up over 70% of isolates from several countries ^12^. More recently, genetic analysis of highly variable single genes for outer membrane proteins such as the 56kDa and 47kDa antigens or more conserved genes (e.g. GroEL) have been used to define genotypic variation. A recent detailed analysis of 56kDa sequences from across South and East Asia identified at least 17 clusters of genotypes belonging to 5 identifiable groups ^13^.

Several multi-locus sequence typing (MLST) schemes using sets of housekeeping genes have been proposed, though no single scheme has been universally accepted ^14–18^. Using one MLST scheme, human isolates from 3 regions of Laos and an isolate from nearby Udon Thani in Northeast Thailand were compared. Low levels of population differentiation were reported between geographically close (Vientiane and Udon Thani) strains, while isolates from southern Laos formed a distinct population ^16^. In that study, 8% of isolates appeared to represent mixed infection, and in Thailand 25% of infections were reportedly mixed ^18^. Recent whole-genome phylogenetic comparisons between 8 well-characterised strains revealed relationships that were significantly different from phylogenies created from single-gene or MLST schemes, illustrating the increased resolution achievable from whole-genome sequencing ^19^. At the level of individual genes such as 56kDa, enormous genetic variability is seen, while at the MLST level only a few clonal clusters are evident.

Several factors combine to make genomic studies of *Orientia* infection challenging. The bacterium is an obligate intracellular pathogen, necessitating cell culture for laboratory propagation ^20^. *Orientia* is typically collected from a range of specimen types including human whole blood, buffy coat and eschar tissue, rodent blood and organs, and chiggers, and the absolute quantity of *O. tsutsugamushi* DNA present in these specimen types is variable, but frequently low. *Orientia* can only be propagated in cell culture, which is technically demanding ^20^ and costly and must be performed in biosafety level 3 facilities ^21^. In one study of 155 infected human blood samples tested by 16S PCR, the median pathogen genome load was 0.013 copies/μL, the interquartile range 0-0.334 and the maximum 310 ^22^, while a recent study from Thailand reported a range of 13.8 to 2,252 copies/μL ^23^. Very few data are available for the quantity of *O. tsutsugamushi* in individual chiggers and there are no published data from rodents. The *O. tsutsugamushi* genome is relatively poorly defined, with just nine complete genome sequences, and shows a high density of repetitive elements and extreme rates of genomic rearrangement, two added challenges that make innovative approaches to sample preparation, sequencing and analysis essential^19,24,25^.

Next-generation sequencing (NGS) techniques have become the gold standard for revealing the genetic variation of organisms ^26^. Culture of *O. tsutsugamushi* in eukaryotic cells can increase the quantity and concentration of DNA available for downstream whole-genome sequencing by thousands of fold. This technique is technically demanding, costly, time-consuming and prone to contamination. Handling infected-cell cultures is also hazardous and carries a risk of infection in those accidentally exposed ^21^. The entire process must be undertaken in biosafety level 3, with all its associated costs and complications.

Targeted enrichment sequencing is a tool whereby certain pre-selected regions of the genome are targeted for sequencing, via hybridisation to a set of probes corresponding to the sequences of interest. The method is akin to, and works similarly to, whole-exome sequencing where just the “exome” or coding portion of the human genome is sequenced. Targeted enrichment can be useful where the whole genome is not required, or a particular genome of interest is selected from contaminating DNA ^27,28^, for example in the metagenomic analysis of multiple virus species, where culture is difficult and costly ^29–31^, and for *Neisseria meningitidis* directly from cerebrospinal fluid, where culture often fails due to prior antibiotic treatment ^32^. Thus, the method in principle provides an efficient alternative to cell culture combined with whole-genome sequencing for *Orientia.*

In summary, the many difficulties associated with conducting a large-scale study at the whole-genome level of *O. tsutsugamushi* in human, chiggers and small mammal samples prompted the development of a probe-based targeted enrichment sequencing strategy, which was used to examine phylogeographical relatedness of samples collecting in Northern Thailand and elsewhere.

## Materials and Methods

### Sample collection

Small mammals were trapped alive in wire-mesh traps baited with corn. Animals were killed using the inhalational anaesthetic isoflurane. Chiggers were collected from rodents by removing the ears and placing into tubes containing 70% ethanol and stored at 4**°**C. The rodent lung, liver and spleen were removed, preserved in 70% ethanol and stored at −80**°**C ^33^. International standards were stringently followed for animal-handling and euthanasia procedures ^34,35^. Free-living chiggers were collected using the black plate method ^36,37^. Human blood and eschar samples were collected during the non-malarial fever studies in Laos ^16^ and the natural immune response to paediatric scrub typhus study in Thailand and stored at −80**°**C ^38^. Chiggers were identified using autofluorescence and bright-field microscopy ^39^ with reference to a range of taxonomic keys ^40–42^. Ethical approval was obtained from Kasetsart University Animal Ethics Committee (EC), Bangkok, Thailand for animal collection; the Faculty of Tropical Medicine EC, Mahidol University, Bangkok, the Chiangrai Prachanukroh Hospital EC, the Chiangrai Provincial Public Health EC and the Oxford Tropical Research EC for human samples in Thailand and additionally the Lao National Committee for Health Research for human samples in Laos.

### DNA extraction and PCR

DNA was extracted from individual chiggers, pools of chiggers, rodent tissues and human samples using the Qiagen Blood and Tissue Kit (Qiagen, USA). The procedures prior to protein digestion were as follows. Chiggers were rinsed with distilled water and individuals cut through the mid-gut using a sterile 30G needle under a dissecting microscope and pools crushed using a sterile polypropylene motorized pestle (Motorized pellet pestle Z35991, Sigma Aldrich, St Louis, MO). Rodent tissues were cut into a small piece (≤10mg of spleen or ≤25mg of liver or lung). Buffy coat or whole blood was extracted from a starting volume of 200 µl. Eschars were collected either as pieces of crust in 70% ethanol or swabs. Chigger, rodent and eschar swabs were incubated with proteinase K at 56°C for 3 hours. Whole blood and buffy coat was incubated for 1 hour and eschar crust was incubated overnight. The rest of the steps followed the manufacturer’s protocol. Chigger samples were eluted in 45 µl, while rodent and human samples were eluted in 100 µl of buffer AE (Qiagen, Hilden, Germany). Samples were stored at −20°C before PCR.

Quantitative real-time PCR targeting the 47kDa *O. tsutsugamushi* outer-membrane protein was performed on all rodent, chigger and human samples ^43^. A PCR master mix was prepared by combining the following reagent volumes per sample: 15 µl of Platinum PCR Supermix UDG (Sigma Aldrich, USA), 0.25 µl each of Forward and Reverse Primers (10 µM) and 0.5 µl of Probe (10 µM). For chigger samples 4 µl of sterile water and 5 µl of DNA was added. For rodent and human samples 8 µl of sterile water and 1 µl of DNA added to complete the Master Mix. PCR was run with the following conditions: 2 minutes at 50°C, then denaturation at 95°C for 2 minutes, followed by 45 cycles of 95°C for 15 seconds and 60°C for 30 seconds. Real-time PCR was performed on a Bio Rad CFX96 (Bio Rad, USA) using in-house quantitative standards. Duplicate 10-fold concentrations from 10^0^ to 10^6^ (1 µl each) and two no-template controls were included on every run.

### Library preparation

In the first round of sequencing in this study, the Nextera XT DNA library preparation kit (Illumina Inc, San Diego, USA) methodology was used to prepare libraries, predominantly for human-derived samples. High duplication rates and relatively low coverage for this approach resulted in a switch to a whole-genome amplification (WGA) step prior to a ligation-based library preparation method.

For Nextera XT libraries, DNA was normalized for an input of ≤1 ng in 5 μL across all samples and libraries were prepared following the manufacturer’s protocol.

For whole-genome amplified libraries, specimens from input volumes ranging from 40 μL (chiggers) and ~50 μL (human samples), to 95 μL for small mammal samples were dried using a Speed-Vac (Eppendorf, Hamburg, Germany) and resuspended in 2.5 μL of TE. WGA was performed following the manufacturer’s protocol for the REPLI-g Single Cell Kit (Qiagen, Hilden, Germany).

The concentration of the amplified DNA was assessed using a Qubit dsDNA HS Assay (Thermo Fisher, MA, USA). Samples were normalized to 500 ng mass in 34 μL DNA and fragmented using an Episonic instrument, (EpiGentek, NY, USA) with the following settings: Amplitude 40, Process time 00:03:20, Pulse-ON time 00:00:20, Pulse-OFF time 00:00:20. The fragmented DNA was cleaned with a 1X ratio of AMPure XP beads (Beckman Coulter, Indianapolis, USA), resuspended in 34 μl.

Libraries were prepared using the NEBNext Ultra DNA Library Prep Kit for Illumina (New England Bioalabs) with a modified protocol. In detail, 6.5 μL NEBNext End repair reaction buffer, 0.75 μL NEBNext End prep enzyme mix and 24.25 μL nuclease-free water were added to each sample and incubated at 20°C for 30 mins and 65°C for 30 minutes. Next, ligation of an in-house Y-adapter was performed by adding 3.75 μL of Blunt/TA Ligase master mix, 1 μL of Ligation enhancer, 1.5 μL of 15 μM adapter and 12.25 μL of nuclease-free water to each sample. This was then incubated for 15 minutes at 20°C, followed by an AMPure XP bead clean-up using 86.5 μL of beads and finally eluted into 100 μL EB buffer.

For sequencing on the Illumina HiSeq4000, an AMPure XP size-selection was then performed by adding 52 μL of AMPure XP to the DNA, mixing, incubating for 5 minutes at room temperature and then transferring to a magnet for 8 minutes. The supernatant was then transferred to a fresh plate and the process repeated using 25 μL of AMPure XP. Finally, the beads were washed twice with ethanol and resuspended in 20 μL of EB buffer.

PCR was then performed on the library using 10 μL of Pre-PCR library, 5 μL of indexed primer i5 and i7, 10 μL water and 25 μL NEBNext Q5 PCR Master Mix. The following conditions were used: 98°C for 30secs, 98°C for 10secs, 65°C for 30secs, 72°C for 30secs, 72°C for 5mins and 10 cycles performed.

A final AMPure XP bead clean-up was carried out using 37.5 μL of beads and eluted in 30 μL of EB buffer. Qubit and Tapestation DNA analysis was performed for all libraries prior to target enrichment.

### Target enrichment

Paired-end DNA libraries prepared using either WGA followed by an in-house library preparation, or Nextera XT, were pooled for capture using pre-designed Agilent SureSelectXT Custom 3-5.9Mb probes and the capture module of the SureSelectXT Reagent Kit, HSQ (Agilent).

The pool of indexed libraries was first normalized to 750 ng in 3.4 μL. A Master Mix containing 2.5 μL of SureSelect Indexing Block #1, 2.5 μL SureSelect Block #2, 3 μL IDT xGen Blocking Oligos was prepared. This was added to the sample, mixed and placed on a thermocycler at 95°C for 5 minutes and then 65°C for 5 minutes.

Next the Hybridization Buffer Master Mix (SureSelect Hyb #1 to #4 and RNase Block) in a total volume 13.5 μL was prepared. 5 μL of baits were aliquoted and added to the Hybridization Buffer Master Mix. This was then transferred to the samples held at 65°C and incubated for 24hrs.

Dynabeads MyOne Streptavidin T1 beads were prepared using the manufacturer’s standard protocol. The PCR plate was maintained at 65°C while moving the samples to the bead plate and pipette mixing. Samples were then incubated on a mixer at 1100 rpm for 30 minutes at room temperature. Samples were then spun briefly, place on a magnetic rack and the supernatant removed and saved. The beads were resuspended in 200 μL of SureSelect Wash Buffer 1 and incubated for 15 minutes at room temperature, replaced on the magnetic rack and the supernatant discarded. The procedure was repeated with SureSelect Wash Buffer 2, incubated for 10 minutes at 65°C and discarding the supernatant as before. The process was repeated 3 times. The beads were then resuspended in 30 μL of distilled water, of which 14 μL was transferred to a post-hybridization PCR using the following PCR Master Mix (Herculase II Reaction buffer, 100mM dNTP Mix, qPCR Library Quantification Primer Premix, nuclease free water and Herculase II Fusion DNA Polymerase), with the cycle parameters of: 98°C for 2mins then 14 cycles of 98°C for 30secs, 57°C for 30secs, 72°C for 1 min, followed by a final extension of 72°C for 10 minutes.

### Sequencing

Sequencing was performed on the Illumina HiSeq4000 with paired-end 150 bp reads.

### Bioinformatic analysis

Raw reads generated from Illumina HiSeq4000 were mapped to the UT76 reference genome (GCF_900327255.1) using BWA MEM v0.7.12 ^44^. Samtools flagstat v1.8 was used to summarise the total number of reads and the proportion mapping to the reference. The reads were then deduplicated using Picard MarkDuplicates v2.0.1 and the same statistics were recalculated, along with the total number of fragments present in the library. Depth of coverage across the whole genome and the proportion of the core genome represented at 1x, 5x and 10x minimum per-base coverage was calculated using GATK v3.7 ^45^.

Haploid variant calling and core genome alignment was performed using Snippy v4.3.6 ^46^. The method identified single nucleotide polymorphisms (SNPs) between the sequence reads and the reference genome. The variant calls were used as input to construct maximum-likelihood (ML) phylogenetic trees using iqtree v1.3.11 ^47^. The most suitable model was selected using ModelFinder Plus which computes the log-likelihoods of an initial parsimony tree for many different models and the Akaike information criterion (AIC), corrected AIC and Bayesian information criterion (BIC) ^48^. To estimate branch supports of the phylogenetic tree inferred from the multiple sequence alignment, ultrafast bootstrap approximation was used ^49^.

### Data availability

The sequences uploaded to generate Agilent SureSelect capture probes are available through Figshare at 10.6084/m9.figshare.12546377. The sequence reads are available in the Sequence Read Archive under project PRJEB39975. For sequence read sets obtained from human samples, reads mapping to the human genome using Bowtie2 were removed from the data before uploading.

## Results

A total of 184 small mammals were trapped at 5 sites in Northern Thailand: Ban Thoet Thai (20.24**°**N, 99.64**°**E), Mae Fahluang district; Ban Song Kwair (20.02**°**N, 99.75**°**E) and Ban Mae Khao Tom (20.04**°**N, 99.95**°**E) and Ban Mae Mon (19.85**°**N, 99.61**°**E), Meuang district in Chiang Rai Province and Ban Huay Muang (19.14**°**N, 100.72**°**E), Tha Wang Pha district, Nan Province. One chigger sample was collected on the Penghu Islands, Taiwan (23.57**°**N, 119.64**°**E). Human samples were collected from Chiang Rai Province, Northern Thailand, across Laos and one from Green Island, Taiwan (22.66**°**N, 121.49**°**E).

### Probe design

The probes were designed in the following way, aiming to ensure that the full diversity of the *O. tsutsugamushi* genome would be successfully captured. Two finished reference strains (Boryong and Ikeda) plus seven other assemblies available at the time of probe design were used (Gilliam: GCF_000964615.1, Karp: GCF_000964585.1, Kato: GCF_000964605.1, TA716: GCF_000964855.1, TA763: GCF_000964825.1, UT144: GCF_000965195.1, UT76: GCF_000964835.1). The complete Boryong strain was used as a reference genome and the whole genome was included in the probe design. To cover genes not found in the Boryong genome, or which had high levels of divergence from the Boryong genome, the genome assemblies were reannotated using Prokka v1.11 and predicted open reading frames from all eight genomes were clustered into groups based on >=80% identity at the protein sequence level using Roary v3.6.0 ^50^. For each cluster, an alignment of the corresponding DNA sequences (using Clustal Omega ^51^) was divided into windows of 120 nt in which every aligned sequence was a candidate probe. Probes were then chosen until every sequence in each cluster was represented by a probe with <10% DNA sequence, a strategy informed by previous work demonstrating efficient capture with probe target divergence up to 20% ^31^ and the requirement to capture as-yet uncharacterised sequences. The reference Boryong gene sequence was always included if it had a representative in the cluster under consideration and sequences that would capture human and rodent genomes (*Rattus norvegicus*) were excluded. The probe design strategy generated a total sequence length of 4.7Mb which was synthesised as a single Agilent SureSelect probe pool. The FASTA file containing the sequences uploaded for probe design is available at 10.6084/m9.figshare.12546377.

### Validation using spiked samples

To create the spike-in solution, DNA was extracted from 20 chiggers of the genus *Walchia* that had previously tested negative for *O. tsutsugamushi* using the 47 kDa real-time PCR. DNA extraction was performed using the methods described previously. The 20 extracted DNA samples (40μL each) of negative chiggers were pooled and then split into 20 tubes, such that the sample was equivalent to the mean amount of DNA extracted from a chigger.

*O. tsutsugamushi* (strains UT76 and CRF136) DNA extracted from cell culture was used to create the dilution series. The concentration was 838 ng/μL with 82% of the DNA being from *O. tsutsugamushi* and 18% from contaminants, (as estimated by qPCR and bulk sequencing of the isolate) giving a starting concentration of *O. tsutsugamushi* of 687 ng/μL. 100,000 copies of *O. tsutsugamushi* = 0.227 ng of DNA. 100,000 copies/μL = 0.42 ng/μL of UT76 stock solution (assuming DNA is 82% *O. tsutsugamushi*). To create a final concentration of 0.42 ng/μL equivalent to 100,000 copies/μL: 2 μL of *O. tsutsugamushi* DNA was added to 18 μL of water, mixed thoroughly and 5 μL of this removed and added to 45 μL of water, mixed again and then 2 μL added to 38 μL water. The following concentrations were made following a dilution series using the prepared *O. tsutsugamushi* and chigger solutions: 100,000, 50,000, 25,000, 10,000, 5,000 and 1,000 copies.

The results of the spiked sample sequencing are shown in Figure 1 and Supplementary Table 1. Total reads of 2.2×10^5^ to 8.5×10^6^ were obtained for each sample, with 32-93% of reads mapping to the target genome. Due to the highly repetitive nature of the *O. tsutsugamushi* genome, which varies hugely between strains, we chose to measure coverage statistics by using coverage across 657 core genes previously identified as present in all samples ^19^, covering 685kb of the 2.2Mb genome. The proportion of the core genome covered with ≤ 10 reads ranged from 14.3 to 99.8. The percentage of reads which were identified as sequencing duplicates ranged from 51 to 66%, with a greater duplication rate in the samples with lower quantities of target DNA, as expected.

**Figure 1.**
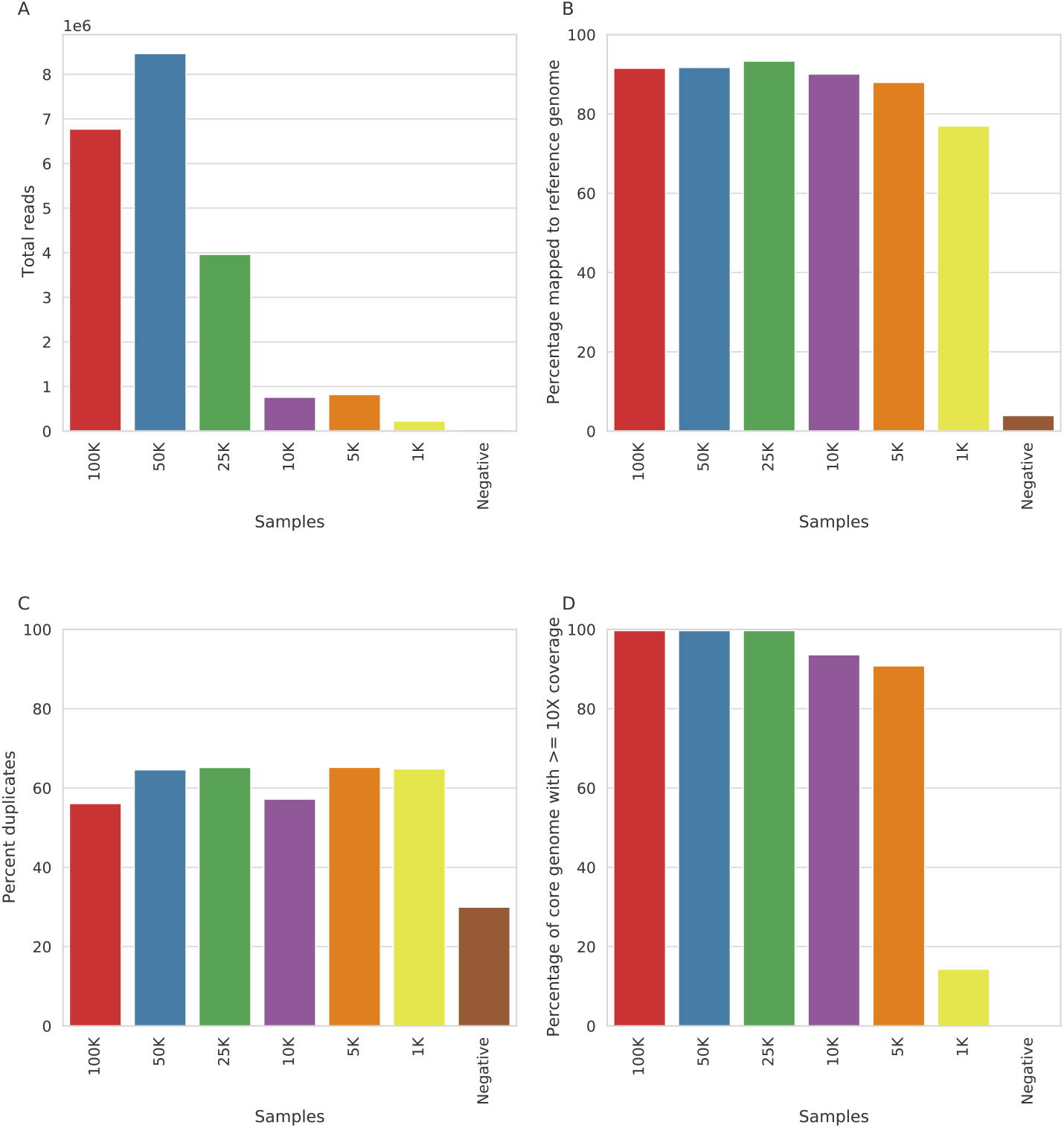
Results from sequencing of spike-in control samples showing a) total reads produced b) percentage of those reads which mapped to the reference genome c) percentage of the reads which were duplicates and d) the percentage of the core genome covered by 10 or more reads.

### Validation of real samples

The low-input Nextera library preparation method was subsequently applied to human samples. This provided inconsistent results, thought to be driven by low and inconsistent amounts of input DNA leading to low-complexity libraries, highly variable pooling and high duplication rates. We therefore altered the library preparation to include a whole-genome amplification step and re-validated using spiked samples, which resulted in lower duplication rates (Supplementary Figure 1 and Supplementary Table 1); all subsequent batches were sequenced with an initial whole-genome amplification step.

Sixty-nine human *O. tsutsugamushi* PCR positive samples from scrub typhus patients were selected from retrospective collections, covering a wide geographical range: 33 from Chiang Rai province in Thailand, 39 from Laos and 1 from Taiwan (Figure 2). Among these, 31 were buffy coat samples, 18 whole blood and 20 eschars (including eschar tissue and eschar swabs). The samples include 11 paired samples with whole-blood/buffy coat and eschar samples from patients collected in Chiang Rai (9 pairs) and Laos (2 pairs).

**Figure 2.**
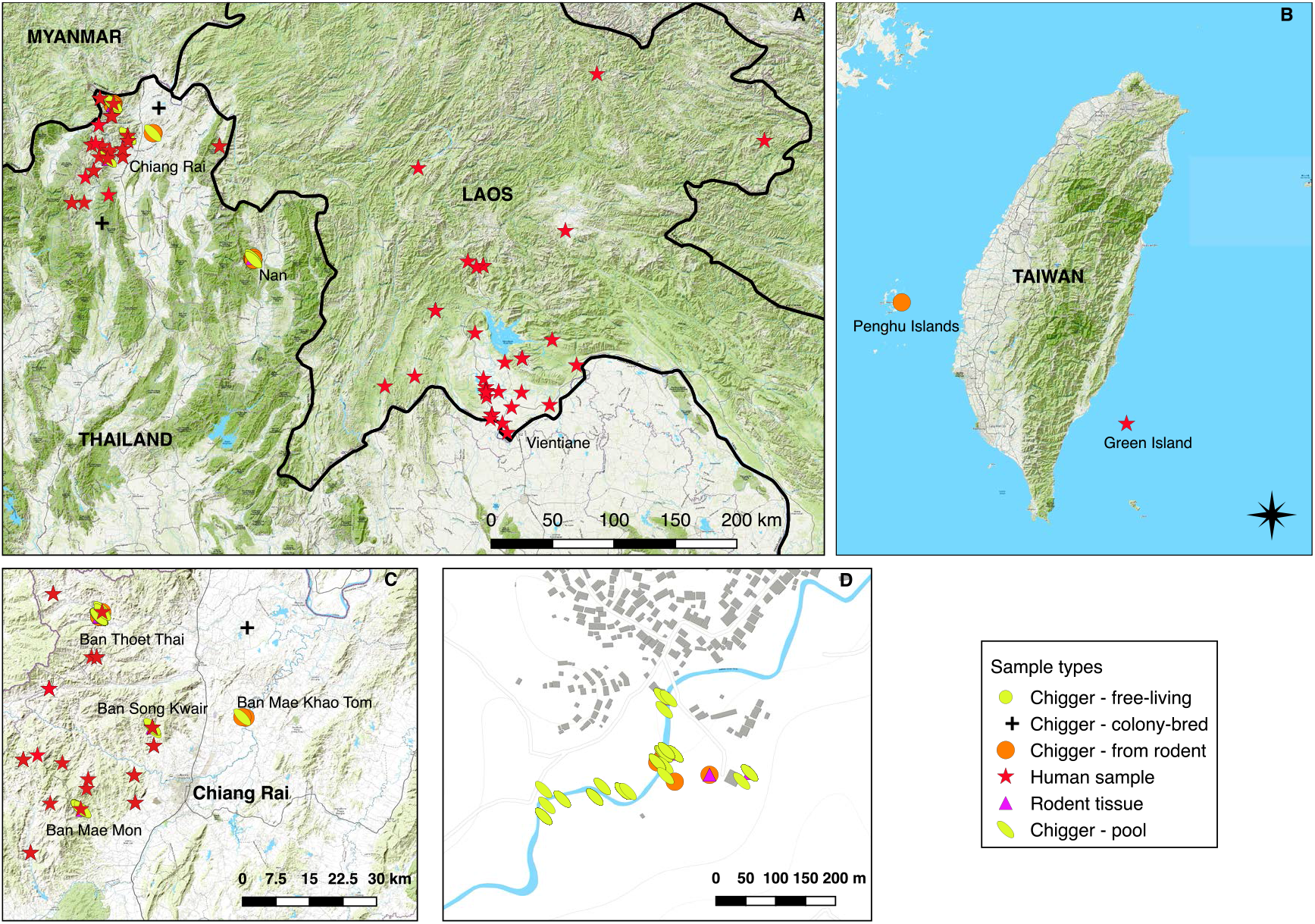
Sample collection locations. A) Southeast Asia with locations in Laos and Northern Thailand, B) Taiwan, C) Chiang Rai Province, with key field sites named, D) Ban Thoet Thai, Chiang Rai Province, site of the greatest number of *O. tsustsugamushi* PCR positive chigger and rodent samples.

Ninety-one *O. tsutsugamushi* PCR positive pooled chigger samples (mean 26 individuals per pool) were selected. These were composed of both pure and mixed species pools collected from 36 small mammals, with multiple pools from some animals (Supplementary Table 1). A total of 27 *O. tsutsugamushi* PCR positive individual chiggers collected from rodents were included of 8 species in 5 genera. These included *L. deliense*, *L. imphalum, Walchia kritochaeta* and *W. micropelta*. Chiggers were collected from 5 sites in Northern Thailand and the Penghu Islands, Taiwan. A single free-living chigger (*L. imphalum*) from Ban Thoet Thai was included. *O. tsutsugamushi*-infected colony chiggers from 3 different species were included, provided by the Armed Forces Research Institute for Medicine (AFRIMS) in Bangkok, Thailand. Six lung and 3 liver tissue samples were included from 7 small mammals of 3 species. Both liver and lung from the same animal were tested in 2 cases. These were collected from 4 sites in Chiang Rai Province (Figure 2).

All samples were PCR positive for the 47kDa gene. The C_t_ values for the samples ranged from 24.6 to 41.3 cycles.

We assessed the sample sequencing based on the number and proportion of reads generated which map to the reference genome, the coverage of the core genes, and the sequence duplication rate (Figure 3). In most samples, only a small proportion of reads mapped to the reference genome, reflecting the performance of the methodology on samples that in general had very small amounts of *O. tsutsugamushi* sequences. Among the different chigger sample types, colony chiggers performed well, with a high percentage of reads mapped to the reference genome likely reflecting their higher input total copy number and corresponding lower C_t_ (mean 29.4, range 28.6-30.2). Chigger pools and individual chiggers from rodents had high variability but with some samples having high levels of reads mapped to the reference genome and correspondingly a high percentage of the genome covered at 10X coverage. C_t_ values for individual chiggers were higher (mean 36.4, median 37, range 30.2-40.2) compared to chigger pools (mean 31.3, median 30.9, range 24.6-40.3).

**Figure 3.**
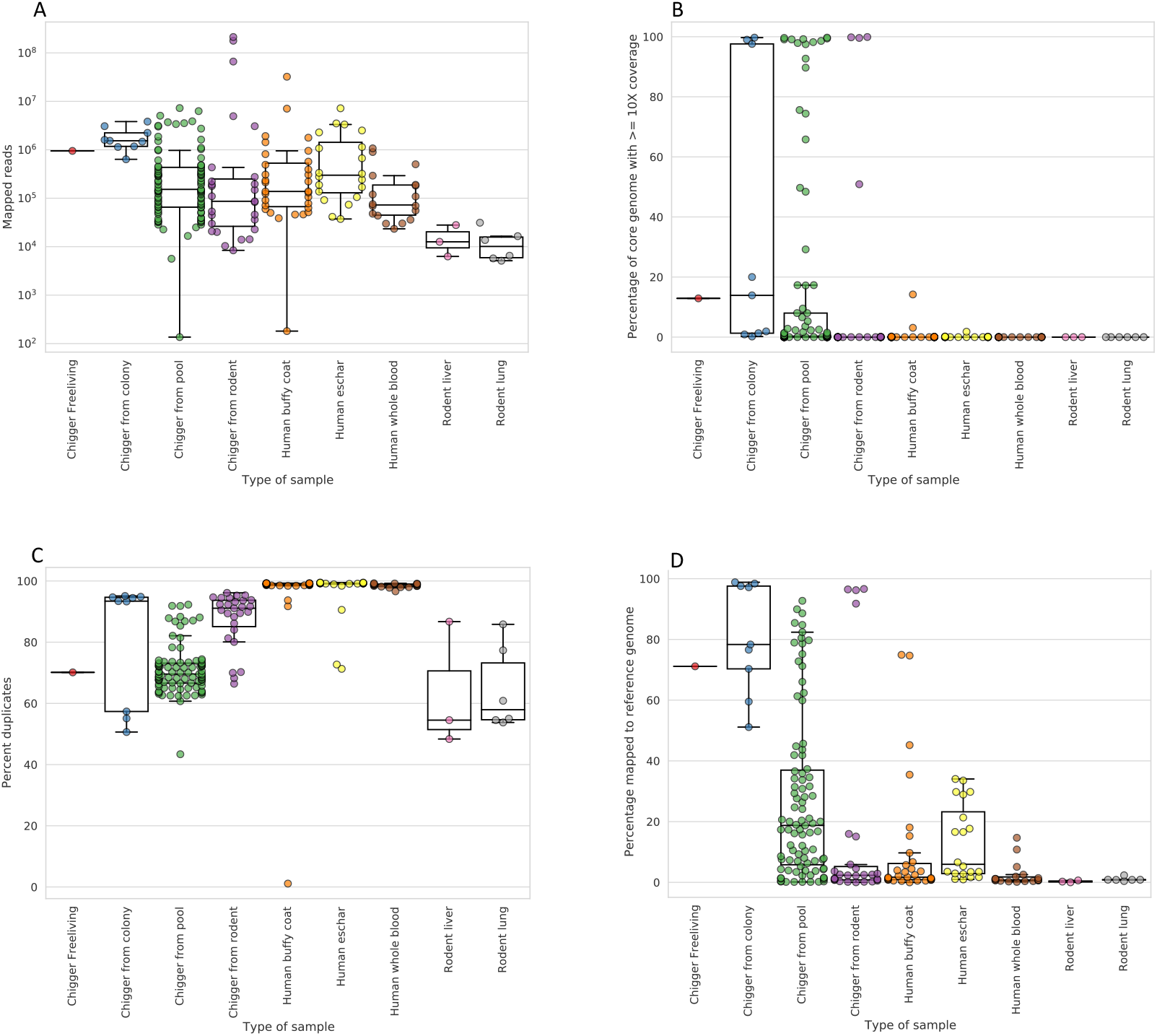
Sequencing statistics for human, chigger, and rodent samples. Panels show a) total number of reads and b) the percentage of reads which were mapped to the reference genome. Panel c) shows the sequence duplication rate and d) shows the coverage of the core genome.

Among the human samples, buffy coat and eschar samples gave more variable performance, with very few samples having sufficient genome coverage to be used in variant calling, and whole blood performed least well with percentage of the core genome covered at 10X or more under 1% in all samples and median percentage of reads mapped to the reference genome of 0.72%. Rodent tissue samples performed poorly in all cases. The relatively low C_t_ values for colony chiggers and their high core genome coverage may reflect the unusual ecological scenario of long-term colony chiggers that may result in higher loads of *O. tsutsugamushi* than wild chiggers.

We expected a positive association between the rate of reads matching *Orientia* sequences and the number of *Orientia* genome copies detectable by qPCR. We compared the fraction of reads which mapped to the C_t_ values (Supplementary Figure 2). Colony chiggers had the highest fraction of reads mapped to the reference genome and tended to have the lowest C_t_ (Supplementary Figure 2). A lower C_t_ (higher input number of genomes) was correlated with the percentage of reads mapped to the reference (Spearman’s rank order correlation=-0.70, p=1.05×10^−35^) (Supplementary Figure 2).

The multiple sample types had a wide range of estimated genome copies, as well as different properties such as total DNA content, which change the ratio of target to non-target DNA. Many samples fell near the lower limit of detection of the qPCR assay, with 69/205 (34%) had a C_t_ value of >35 It appears that a C_t_ of ≥35 results in poor coverage and low percentage mapping to the reference.

Variant calling was performed on the entire set of sequenced samples. Due to the low sequence coverage for many samples, phylogenetic comparisons were attempted only for a set of 31 samples with >50,000 bases called: 4 chigger pools from Ban Mae Mon, Thailand, 1 human buffy coat sample from Na Meuang, Laos, 1 individual chigger from the Penghu Islands, Taiwan, 4 individual chiggers and 17 chigger pools from Ban Thoet Thai, Thailand, and 4 colony chiggers. The median C_t_ value for these samples was 29.0 (range 25.4-34.2). The distribution of bases called for these 31 samples is shown in Supplementary Figure 4. Coverage for each core gene is shown in the heatmap in Supplementary Figure 5. For almost all samples, there is some sequence coverage for each of the core genes, and for those with fewer positions called it is due to incomplete coverage across the genome rather than genes which are completely uncovered in sequencing. A notable exception is sample C0546, which has many genes which have no coverage at all but sufficient coverage in the remaining genes to meet the 50,000bp threshold. A small number of genes were completely uncovered in multiple samples, most notably several genes which have no coverage in any of the samples taken from the R240 pools from a rodent in Ban Mae Mon.

**Figure 4:**
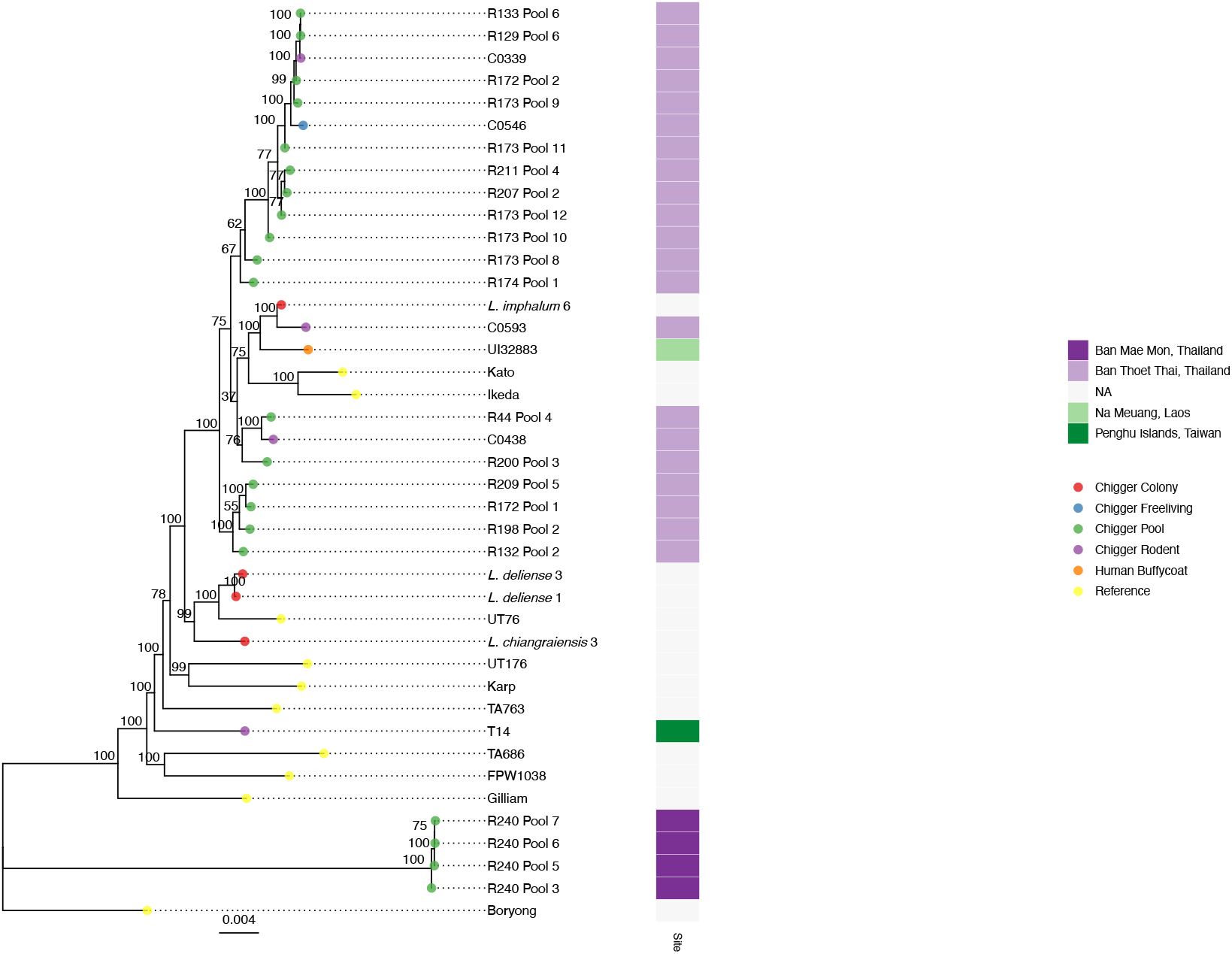
A maximum-likelihood phylogenetic tree produced using IQTREE from all samples that have >50kb of called positions. Tip colors represent the source of each sample, and the heatmap shows the site where samples were collected. The node labels show ultrafast bootstrap support values.

The phylogeny is show in Figure 4. Branch bootstrap values, which can be interpreted as the relative (%) support of the data for the tree topology represented by the pairings of isolates or groups of isolates on either side of the labelled branch, are plotted on the tree and fall below 70% support for some branches, indicating some uncertainty in tree topology. The samples include two colony chiggers from the same *L. deliense* colony. These samples are closely related but not identical (35 SNPs between the two samples).

## Discussion

We have successfully developed and tested the first whole-genome sequencing of *O. tsutsugamushi* performed without prior cell culture. The sequence data generated provided an opportunity to compare *O. tsutsugamushi* strains with greater resolution than previously possible.

The sequencing results displayed great variability, with sufficient success to call variants and perform phylogenetic analysis in a proportion of samples from individual chiggers and chigger pools. The yield of unique on-target reads, particularly at the low copy number dilutions (5,000 and 10,000 copies) was higher for WGA before library preparation than for Nextera XT, and the duplication rate was also improved. The low success rate likely reflects very low quantities of *O. tsutsugamushi* DNA present in many samples, especially human samples, and reflects the current limit of our enrichment method which cannot enrich sufficiently to overcome the low levels of input DNA. While no firm C_t_ cutoff value can be established above which target enrichment sequencing cannot be successfully performed, samples with a C_t_ value of 35 or less are candidates for sequencing. Methods for human and rodent DNA depletion prior to sequence capture may improve the performance of enrichment. The first full genome of *L. deliense* has been published since this array was designed, and this could be used to check for any sequences in the array design which may capture off-target chigger DNA ^52^.

A recent study has reported phylogenetic comparisons of *O. tsutsugamushi* strains from chiggers collected from the same host animal, based on sequencing of a single gene (encoding the 56 kDa antigen) ^53^. Results revealed mixed infections; with some chiggers containing a single genotype and others mixed genotypes. There is also evidence of different *O. tsutsugamushi* 56 kDa type-specific antigen genotypes being maintained and transmitted transovarially in colony chiggers ^54^.

The sequence capture probes used in this experiment were designed when only two complete genomes were available to use in the design process. Of the incomplete assemblies included in the design process, two strains have been removed from RefSeq due to problems with the assembly, and more complete genomes are now available. A new probe design using the same approach but more genomes may improve the capture efficiency.

Despite the poor performance for the target enrichment sequencing on some samples, we were able to generate a phylogeny using 30 chigger samples, 1 human sample, and 8 complete reference genomes, which represents the first phylogenetic analysis of *O. tsutsugamushi* from chiggers. Among the 31 best-sequenced samples, >98.5% of the core genes of the reference sequence were covered by at least one read at all positions. For most samples, the regions of no coverage were confined to a very few genes, some of which were present in all samples. Intriguingly, for chigger pools from Ban Mae Mon (R240), more genes were incompletely covered, and most of these were present in all samples, even though the total volume of on-target reads (equivalently, the average coverage of the core genome) was similar in these samples as in other high-performing samples. This could be due to diversity in these genes beyond the limits of what our probes are able to capture; however, the sequence capture probes have been shown to be effective at up to 20% sequence divergence ^31^, and the overall diversity between our phylogenetic samples is well below this limit. It is more likely that the set of core genes determined from the known complete genomes is not universally present in all strains.

The study included strains sequenced from chiggers collected from a single host animal, strains from chiggers from several animals at a single study site of <10km^2^ and from two sites 45km apart. Samples from Ban Mae Mon are clearly distinct from samples from Ban Thoet Thai, which group together (Figure 4). All the chigger pools and individuals from Ban Thoet Thai consisted of the known vector *L. imphalum* (with or without some *Walchia* species). The Taiwanese chigger was the known human vector *L. deliense*. The R240 pools from Ban Mae Mon, which form a distinct cluster separate from all other samples, were collected from the scansorial tree shrew *Tupaia glis* and consisted of *L. turdicola* and *Helenicula naresuani* chiggers – neither known to be human vectors nor previously reported as being infected with *O. tsutsugamushi*. The reference genomes, which were collected from five different countries between 1943 and 2010, are spread throughout the tree and many are more closely related to the samples from Ban Thoet Thai than the samples from Ban Mae Mon are to those samples. A possible explanation for this is that *O. tsutsugamushi* has been previously introduced into these two locations from divergent sources and continues to evolve locally on a small scale, and larger-scale *O, tsutsugamushi* movement between locations is a rare event due to the restricted range of the host species.

Important questions remain about the role of recombination between strains in infected chiggers and to what extent the accessory genome of *Orientia* is open or closed. The sequence capture approach used in this study does not recover the complete accessory genome, and hence cannot assist with the latter question.

The accumulation of more high-quality sequences may allow characterization of the recombination landscape. However, *O. tsutsugamushi* genomes are known to have poorly conserved synteny, which is likely to complicate analysis of incomplete genomes.

Among captured sequences, pairwise divergences were in the range of 0-4%, well within the reach of probe-based sequence enrichment for pathogen genomics ^31^. This illustrates the robustness and adaptability of probe-based sequence enrichment, providing a means for genome-wide amplification of sequence information without the need to validate a very large number of PCR primers, any of which could fail because of hitherto uncharacterised sequence variation.

The methods developed in this project have, for the first time in scrub typhus research, demonstrated phylogeographic clustering of *O. tsutsugamushi* strains at international, provincial and highly local scales. This shows that both closely related and more distantly related strains may co-exist in one site. As methods improve and can be applied to a greater range of samples, particularly sympatric rodents and exposed humans, further insights into this fascinating phylogeographic variation will be revealed with important consequences for diagnostic tests and vaccine development strategies.

## Supporting information

Supplementary Figurres

## Acknowledgments

We are very grateful to Associate Professor Bounthaphany Bounxouei, ex-Director of Mahosot Hospital, the Director and staff of the Microbiology Laboratory, LOMWRU and wards, Assistant Professor Chanphomma Vongsamphan, ex-Director of Department of Health Care, Ministry of Health, and H.E. Professor Bounkong Syhavong, Minister of Health, Laos, for their help and support. We thank the Director and staff of the Microbiology Laboratory and the staff of LOMWRU for their wonderful help, Dr Chi-Chien Kuo and colleagues at the National Taiwan Normal University, Taipei for facilitating chigger collection on the Penghu Islands. We thank Sebastiaan Van Hal for providing the human sample from Taiwan. We are very grateful to Rawadee Kumlert at Mahidol University for her assistance in mite morphotyping, and all the staff at the Chiangrai Clinical Research Unit and field teams, in particular Piangnet Jaiboon, Dr Tri Wangrangsimakul, Dr Serge Morand and Dr Kittipong Chaisiri. We thank Prof Alistair Darby for comments on the manuscript.

Material has been reviewed by the Walter Reed Army Institute of Research. There is no objection to its presentation and/or publication. The opinions or assertions contained herein are the private views of the author, and are not to be construed as official, or as reflecting true views of the Department of the Army or the Department of Defense. Research was conducted under an approved animal use protocol in an AAALACi accredited facility in compliance with the Animal Welfare Act and other federal statutes and regulations relating to animals and experiments involving animals and adheres to principles stated in the Guide for the Care and Use of Laboratory Animals, NRC Publication, 2011 edition.

## CReDiT Author statement

**Ivo Elliott:** Conceptualization, Methodology, Formal analysis, Investigation, Writing - Original Draft, Funding acquisition. **Neeranuch Thangnimitchok:** Investigation, Writing – Review & Editing. **Mariateresa de Cesare:** Investigation, Validation, Writing – Review & Editing. **Piyada Linsuwanon:** Resources. **Daniel Paris:** Conceptualization, Methodology, Supervision, Writing – Review & Editing. **Nicholas Day:** Conceptualization, Writing – Review & Editing, Supervision. **Paul Newton:** Conceptualization, Writing – Review & Editing, Supervision. **Rory Bowden:** Conceptualization, Methodology, Formal analysis, Writing – Review & Editing, Supervision, Funding acquisition. **Elizabeth Batty:** Conceptualization, Methodology, Software, Validation, Formal analysis, Data Curation, Writing – Original Draft, Supervision.

