## Supplementary Figurres for "Targeted sequence capture of *Orientia tsutsugamushi* DNA from chiggers and humans"

### Supplementary Figures

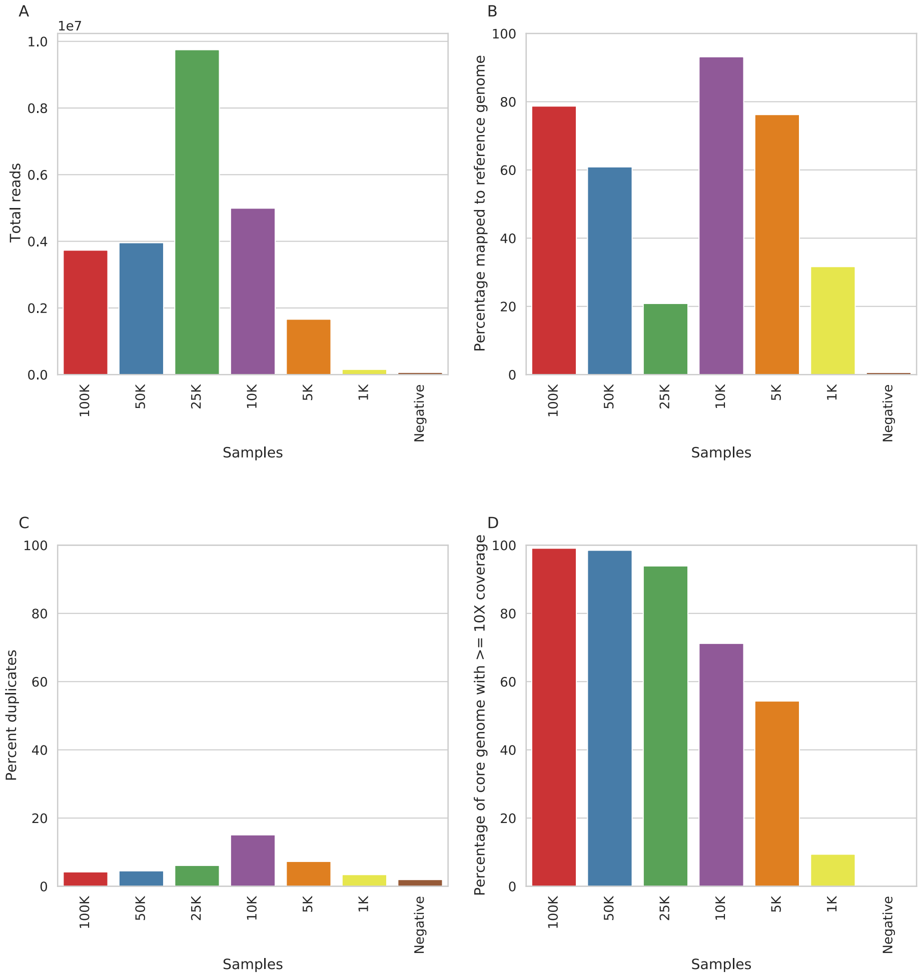


Supplementary Figure 1. Results from sequencing of spike-in control samples prepared using whole-genome amplification methods showing a) total reads produced b) percentage of those reads which mapped to the reference genome c) percentage of the reads which were duplicates and d) the percentage of the core genome covered by 10 or more reads.

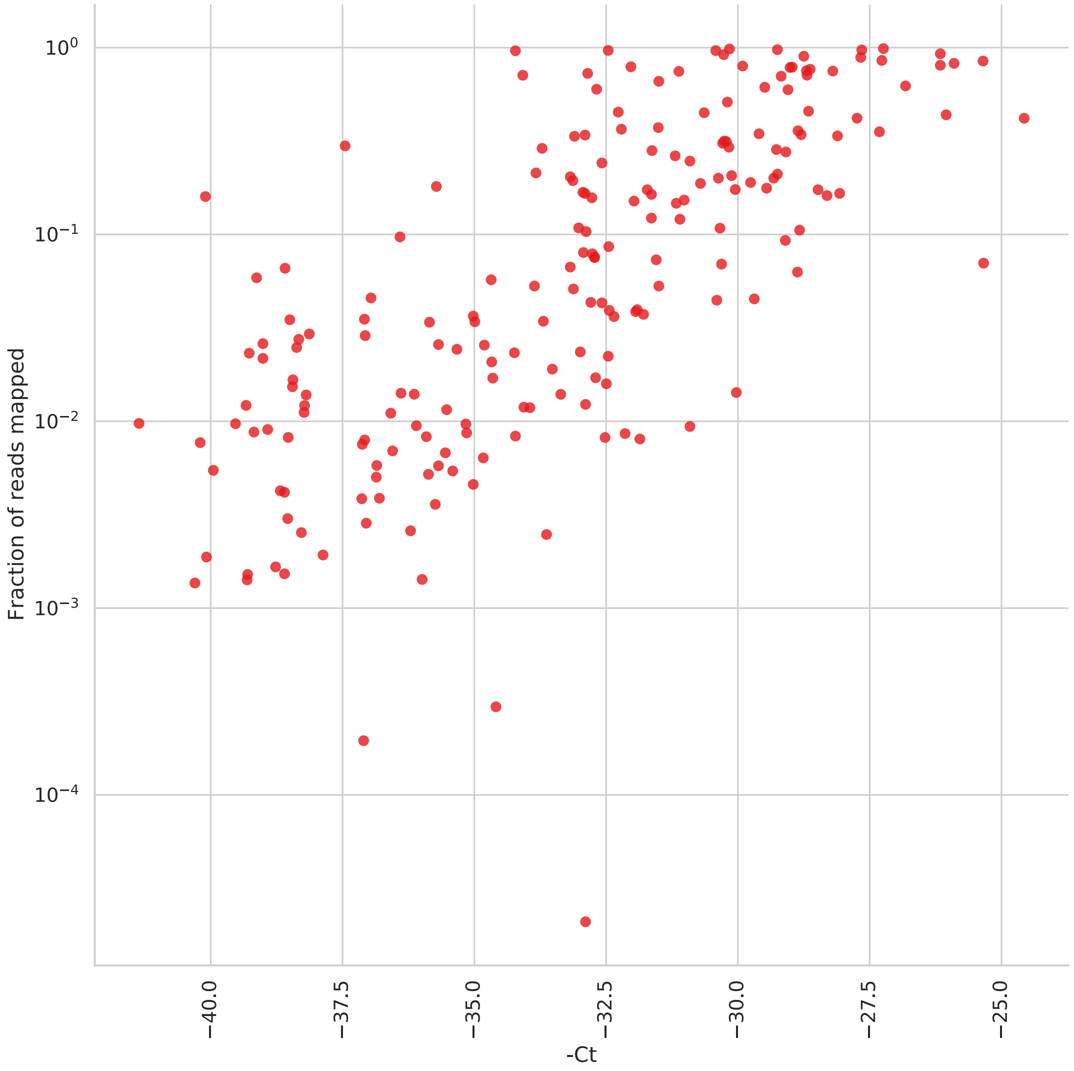


Supplementary Figure 2. Fraction of total reads mapped to the reference genome plotted against - $C_t$.

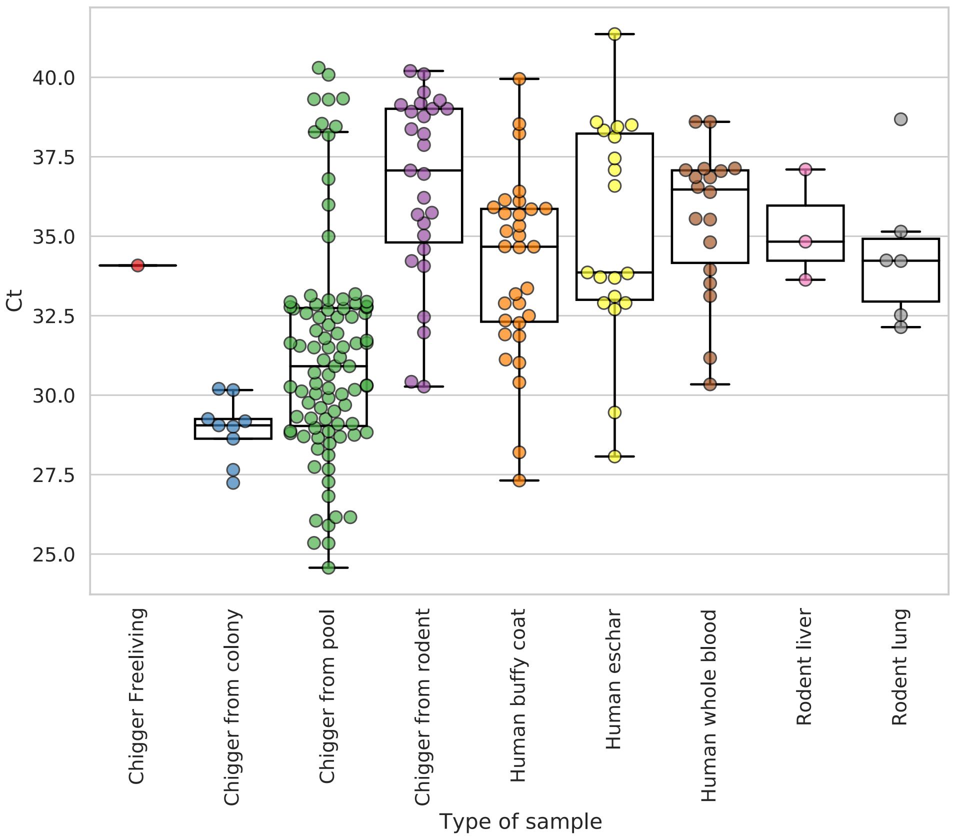


Supplementary Figure 3. $C_t$ values plotted by type of sample.

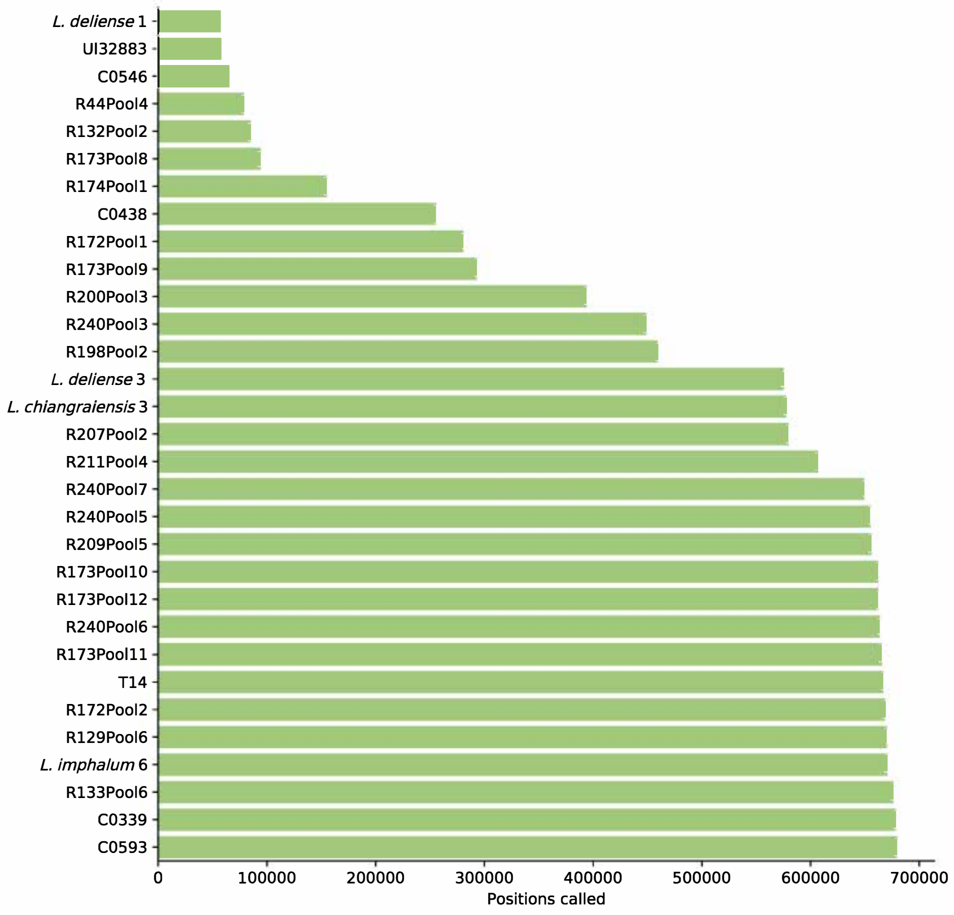


Supplementary Figure 4. Distribution of number of positions called in the 31 samples used for phylogenetic analysis.

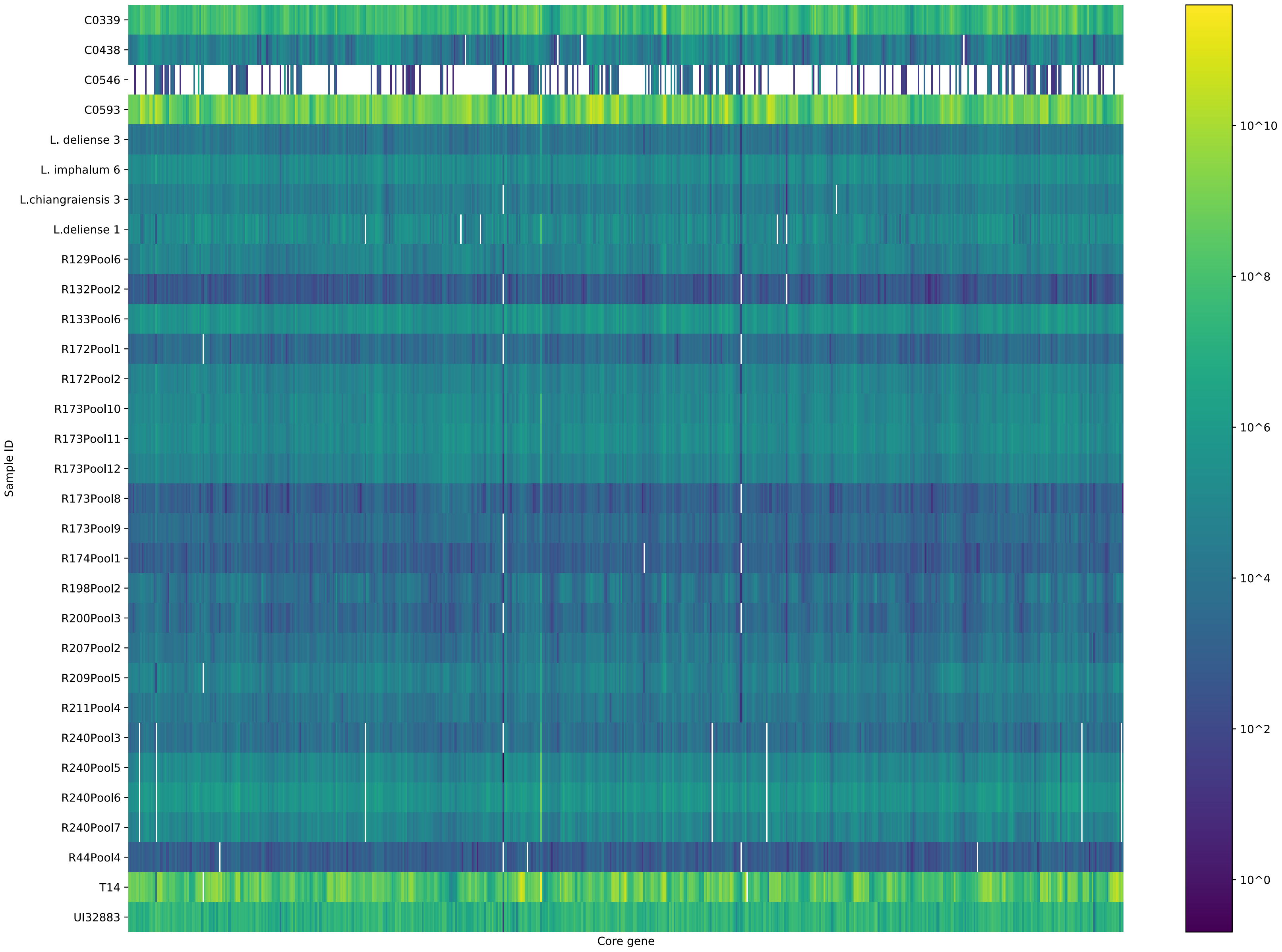

C0339
C0438
C0546
C0593
L. deliense 3
L. imphalum 6
L.chiangraiensis 3
L.deliense 1
R129Pool6
R132Pool2
R133Pool6
R172Pool1
R172Pool2
R173Pool10
R173Pool11
R173Pool12
R173Pool8
R173Pool9
R174Pool1
R198Pool2
R200Pool3
R207Pool2
R209Pool5
R211Pool4
R240Pool3
R240Pool5
R240Pool6
R240Pool7
R44Pool4
T14
UI32883
Sample ID
Core gene
10^10
10^8
10^6
10^4
10^2
10^0

Supplementary Figure 5. Heatmap showing median coverage across each of the 657 core genes for the 31 samples used in phylogenetic analysis. The core genes are plotted in the order found in the UT76 reference genome. Genes with zero coverage are not coloured and appear as missing bars.
